# Ureterorenoscopic (URS) lithotripsy and balloon dilation cause acute kidney injury and distal renal tubule damage

**DOI:** 10.1101/2021.01.12.426334

**Authors:** Ho-Shiang Huang, Chan-Jung Liu

**Affiliations:** Department of Urology, National Cheng Kung University Hospital, College of Medicine, National Cheng Kung University, Tainan, Taiwan

## Abstract

Ureterorenoscopy (URS) is believed to be a safe and effective procedure to treat ureteral stone or ureteral stricture. Rapidly increased intrarenal pressure during URS may have a negative impact on the kidney, but the effect on renal functions is not well known. The aim of this study is to evaluate whether URS balloon dilation or lithotripsy would cause acute kidney injury (AKI), which was evaluated by urine neutrophil gelatinase associated lipocalin (NGAL) and renal tubular damage, which was evaluated by urine α glutathione S-transferase (GST) and πGST. This prospective study consisted of 207 patients with mean age 53.8 years old between September 2012 and June 2013. Four groups were included: the ureteral stricture group (group 1), the ureteral stone group (group 2), and two control groups. URS induced increased uNGAL on both Days 1 and 14 in both groups, and only elevated u-πGST levels were noted on Day 14 after URSL. On Day 14, the difference between low-grade and high-grade hydronephrosis was significant in group 1 (p < 0.001) rather than group 2 (p = 0.150). By multivariate logistic regression analysis, age, baseline eGFR, and stone size > 1.0 cm were associated with complete recovery of hydronephrosis after URS on Day 14. Ureteral stone patients with preserved renal function suffered more AKI (uNGAL) than those with impaired renal function. However, URS-related AKI had no significant difference between stone ≤ 1 cm and > 1 cm subgroups. Besides, urine αGST and πGST were both significantly higher in stone > 1 cm subgroup when compared to ≤ 1 cm subgroup. In conclusion, URS laser lithotripsy and balloon dilatation all resulted in AKI and renal tubular damage on Day 14 though post-URS double-J (DBJ) stenting was performed in every patient.

## Introduction

Impairment of urinary flow due to the urinary tract obstruction, referred to obstructive uropathy, is a manifestation of a variety of kidney and ureteral disease [1]. The progression of renal function after relief of obstructive uropathy has been widely studied. When experimental animals undergo 24 hours (hr) of unilateral ureteral obstruction, a decline in renal hemodynamic and tubular function is found [1, 2]. Glomerular filtration rate (GFR) is directly affected by intrapelvic pressure and will decrease and actually become zero as pressure progressively increases, whereas renal blood flow does not respond directly to intrapelvic pressure [3]. Furthermore, calcium oxalate (CaOx) stone disease *per se* can induce renal tubular damage and renal interstitial fibrosis, which was found both in stone patients and experimental animals [4-6]. Therefore, more profound kidney damage is more likely to be found in patients with obstructive uropathy caused by a CaOx ureteral stone.

Ureterorenoscopic lithotripsy (URSL) is a safe, effective and less invasive method for the treatment of ureteral stones [7]. With advances in technology, ureterorenoscopy (URS) have evolved to a significantly smaller outer diameter. Good irrigation is vital for ureteral dilatation and instrument passage, and irrigation is required to provide a clear vision [8]. However, application of high-pressure irrigation during URS can cause accumulation of renal intrapelvic fluid and increases in intrapelvic pressure significantly [8]. High-pressure irrigation during URS can cause irreversible damage to the urothelium and renal parenchyma [9, 10]. In addition, ureteral stricture is another major cause of obstructive uropathy [11]. The same as ureteral stone, ureteral stricture is also associated with kidney injury and fibrosis [12]. Various managements of ureteral stricture can be used based on urologists’ preference and experience. Balloon dilation of the ureter is a well-accepted surgical technique to resolve ureteral stricture [13]. However, many urologists prefer internal stents, that are double-J stents (DBJ) in most circumstances, to treat ureteral stricture rather than balloon dilation because of the potential risk of ureteral injury. Taken together, although both ureteral stone or ureteral stricture are associated with kidney injury, it is still controversial whether the treatments of them, which are URSL and balloon dilation, attenuate or aggravate the kidney injury. Whether URSL or balloon dilation of the ureteral stricture cause more profound acute kidney injury (AKI) and renal tubular damage has not yet been fully investigated.

Neutrophil gelatinase-associated lipocalin (NGAL) has emerged as the most promising biomarker of AKI [14]. NGAL is a 21-kDa protein expressed in neutrophils and human epithelia and has a physiological role in iron transport, while regulating cell growth and differentiation [15]. Under various types of AKI, NGAL is secreted into urine from the ascending limb of the loop of Henle and the distal nephrons. Growing evidence has suggested that urine NGAL is an early and accurate biomarker to predict AKI [14, 16]. As in our previous studies, kidney stone disease is associated with renal tubule damage as well as lipid peroxidation [17, 18]. α-glutathione S-transferase (αGST) is a cytosolic enzyme that has been proven to be a useful marker of chemically-induced tubular damage, particularly in the S3 segment of the proximal tubule [19]. πGST is also a cytosolic enzyme and is mainly localized in the distal tubules and collecting ducts, and its presence in the proximal tubules is sparse [18].

In this prospective study, we investigated whether URSL or URS balloon dilation would cause AKI (evaluated by urine NGAL level) and renal tubular damage (evaluated by urine αGST and πGST levels). We also evaluated the variables that contribute to complete recovery of hydronephrosis after URS surgery and the impact of hydronephrosis.

## Materials and methods

The protocol of this prospective study was approved by the Institutional Ethics Review Board of National Taiwan University Hospital (Registry Number: 201205117RIC), and all subjects provided written informed consent.

### Study design and populations

We prospectively enrolled patients from a single tertiary medical center between September 2012 and June 2013. All patients were admitted for URS holmium: yttrium-aluminum-garnet (Ho: YAG) laser lithotripsy or balloon dilation because of diagnosed with acute unilateral hydronephrosis caused by ureteral stones or ureteral stricture, which was confirmed by renal ultrasonography, intravenous urography, or non-contrast computed tomography (CT). The exclusion criteria included patients with (1) previous urolithiasis history; (2) acute pyelonephritis or associated urinary tract infection; (3) pre-URSL nephrostomy tube insertion or currently indwelled; (4) hydronephrosis caused by infravesical obstruction, uterine myoma, malignancy, or other retroperitoneal etiology; (5) stone not containing any amount of CaOx based on the stone analysis; (6) other inflammatory or malignant diseases.

### Hydronephrosis classification

The grade of hydronephrosis was classified according to Goertz JK and Lotterman S [22]: grade 1 was defined as enlargement of the calices with preservation of the renal papillae; grade 2 was defined as rounding of the calices with obliteration of the renal papillae; grade 3 was defined as caliceal ballooning with cortical thinning. All renal sonographic examinations and determination of the grade of hydronephrosis were performed by one urologist. The timing of post URS sonography was two weeks after surgery and after DBJ removal.

### Patient grouping

In current study, all subjects were categorized into four groups: two study groups (ureter stone group and ureter stricture group) and two control groups (positive and negative). The diagnosis of obstructive uropathy was established via intravenous urography (IVU) or non-contrast CT scan 2-3 weeks after their first outpatient department interviews. One week after the IVU or CT scan, patients returned to our outpatient department to confirm that the obstructive uropathy was associated with ureteral stones or ureteral stricture. All patients were enrolled into the ureteral stricture group (group 1) or ureteral stone group (group 2) based on the image results.

Serum and 24-hour urine samples were collected at three time periods: Pre-URS (baseline) sample was collected after overnight fasting, 1-day post-URS (Day 1) samples, and 2-week post-URSL (Day 14) samples were collected from all patients while they followed a normal diet. There was at least a 7-10 day interval between IVU examination and subsequent collection of blood and urine samples. The maximal stone length was assessed based on the images of IVU or abdominal CT.

In current study, we used two groups of controls: negative control (NC) and positive control (PC). The NC, serving as the controls at the baseline status, were those who had unilateral ureteral stricture history with long-term unilateral DBJ catheter indwelling. A total of 14 patients were replaced with a 7 Fr. DBJ catheter under anesthesia, and there was no recurrence of ureteral stricture confirmed by URS and retrograde pyeloureterography at the same time. We used patients with unilateral renal staghorn stone and receiving percutaneous nephrolithotomy (PCNL) as the PC, and they served as the controls mainly at the post-URS period to investigate the impact of URS on the changes of these biomarkers. Before PCNL, the PC received URS with a 7 Fr. ureteral DBJ indwelling. The NC represented the patients without current evidence of obstructive uropathy.

Otherwise, the PC represented those who got most severe renal injury because of inevitable renal volume damage during PCNL.

### The procedure of ureterorenoscopy

In stone patients (group 2), the ureteral stone was disintegrated by the application of Ho:YAG laser (Odyssey, Convergent Laser Technologies, Alameda, CA,USA) through URS. After the ureteral stone had been disintegrated, ureteral patency was examined by URS to ensure that there was no ureteral injury caused by laser lithotripsy.

In ureteral stricture patients (group 1), high pressure balloon catheter (UroMAx Ultra, Boston Scientific, Natick, MA, USA) was applied to relieve the stricture. Balloon dilation was applied at least two times with balloon inflation pressure up to 18-20 atmosphere (ATM) for 5 minutes each time under fluoroscopy to ensure all segments of ureteral stricture were relieved.

### Biochemical analysis and renal function determination

Serum creatinine (Cr) level was examined twice, which were the baseline and 2-week post URS, in every enrolled patient and the eGFR was calculated using the formula as 186×(Serum Cr)^-1.154^×(age)^-0.203^×(0.742 if female)×(1.210 if African-American) [2, 3].

Commercial kits were used to determine the urine level of NGAL (NGAL ELISA Kit, BioPorto Diagnostics A/S, Copenhagen, Denmark) and the urine levels of stone-induced renal tubular damage markers; namely, the urinary αGST(Alpha GST EIA, Argutus Medical, Dublin, Ireland), which is a marker of proximal tubular damage, and the πGST (Pi GST EIA, Argutus medical, Dublin, Ireland), which is a marker for distal tubular damage. Urinary αGST and πGST were examined at baseline and 2-week post-URS (Day 14) and all assays were performed in duplicate. Urine NGAL (uNGAL) level was examined at three time periods: baseline, post-URS Day 1 and post-URS Day 14.

### Statistical analysis

Continuous variables are presented as the mean values ± standard deviation, whereas categorical variables are presented as frequencies. Two sample comparisons between kidney stone patients and controls were performed using the unpaired Student’s *t*-test, Mann-Whitney *U*-test or Fisher’s exact probability test, as appropriate. Comparisons across the three groups were done using the *chi* square test for categorical variables and by analysis of variance (ANOVA) or Kruskal–Wallis test for continuous variables depending on the distribution of the variable. Logistic regression was used for the univariate and multivariate analyses to identify factors having an effect on hydronephrosis recovery (hydronephrosis degree = 0). Pearson’s correlation coefficient (γ) was used to assess the correlation between clinical variables. In all tests, *p* value <0.05 was considered statistically significant.

## Results

A total of 220 patients were enrolled in current study and mainly male patients (67.3%) with a mean age of 53.80 (Table 1). There was no significant different in age, eGFR, BMI and 24-hour urine output between these four groups. In group 1, the most common site of ureteral stricture was upper ureter (59.6%); whereas the most common location of ureteral stone was also upper ureter (56.0%). The mean maximal stone length was 0.9 cm.

**Table 1.** Clinical Characteristics of the study population.

|  | Negative control | Ureteral stricture (Group 1) | Ureteral stone (Group 2) | Positive control |
| --- | --- | --- | --- | --- |
| Number | 13 | 52 | 141 | 14 |
| Hydronephrosis (No.) |  |  |  |  |
| Grade 0 | 14 | 13 | 5 | 0 |
| Grade 1 | 0 | 17 | 61 | 11 |
| Grade 2 | 0 | 13 | 58 | 3 |
| Grade 3 | 0 | 9 | 17 | 0 |
| Age, years | $51.7 \pm 2.6$ | $53.0 \pm 2.0$ | $54.5 \pm 1.1$ | $59.1 \pm 2.7$ |
| Male | 8 (61.5) | 28 (53.8) | 101 (71.6) | 11 (78.6) |
| eGFR (mL/min/1.73 m<sup>2</sup>) | $92.0 \pm 6.6$ | $87.0 \pm 4.0$ | $86.9 \pm 3.5$ | $85.1 \pm 8.7$ |
| BMI | $25.9 \pm 1.2$ | $24.6 \pm 0.5$ | $24.3 \pm 2.1$ | $24.5 \pm 1.0$ |
| Stone size (cm) | 0 | 0 | $0.9 \pm 0.5$ | $5.4 \pm 0.7$ |
| Location (stone or |  |  |  |  |
| stricture) |  |  |  |  |
| Upper ureter | - | 31 (59.6) | 79 (56.0) | - |
| Middle ureter | - | 3 (5.8) | 42 (29.8) | - |
| Lower ureter | - | 18 (34.6) | 20 (14.2) | - |

Data expressed as mean ± standard deviation or number (percent).
BMI: body mass index; eGFR: estimated glomerular filtration rate.

### The impact of different degree of hydronephrosis on kidney injury biomarkers

Table 2 shows the comparisons of different kidney injury biomarkers in different degree of hydronephrosis in group 1 and group 2. To compare with the NC group, significantly elevated uNGAL level was noted in stone patients (group 2) with moderate-to-severe hydronephrosis and severe hydronephrosis in ureter stricture patients (group 1). Significant elevation of renal tubular damage markers were also found in stone patients with moderate-to-severe hydronephrosis, but we didn’t find any difference in stricture patients. Interestingly, when compared with stricture patients, the levels of u-αGST and u-πGST were significantly higher in moderate-to-severe hydronephrosis of stone patients; whereas there was no significant difference in uNGAL between groups.

**Table 2.** The impact of hydronephrosis on kidney injury, evaluated by acute kidney injury (AKI) maker (NGAL) and renal tubular damage markers (αGST, πGST) in group 1 (ureteral stricture) and group 2 (ureteral stone) at baseline condition when compared to the negative controls.

**Part 1. Compared with negative controls (NC)**
| **NGAL** | **P value** |  | **P value** |
| --- | --- | --- | --- |
| stone H0 vs. NC | 0.456 |  |  |
| stone H1 vs. NC | 0.153 | stricture H1 vs. NC | 0.113 |
| stone H2 vs. NC | **0.035** | stricture H2 vs. NC | 0.390 |
| stone H3 vs. NC | **0.006** | stricture H3 vs. NC | **0.007** |
| **αGST** |  |  |  |
| stone H0 vs. NC | 0.241 |  |  |
| stone H1 vs. NC | 0.058 | stricture H1 vs. NC | 0.385 |
| stone H2 vs. NC | **0.002** | stricture H2 vs. NC | 0.848 |
| stone H3 vs. NC | **0.000** | stricture H3 vs. NC | 0.352 |
| **πGST** |  |  |  |
| stone H0 vs. NC | 0.332 |  |  |
| stone H1 vs. NC | 0.059 | stricture H1 vs. NC | 0.080 |
| stone H2 vs. NC | **0.007** | stricture H2 vs. NC | 0.604 |
| stone H3 vs. NC | **0.000** | stricture H3 vs. NC | 0.440 |

**Part 2. Compared between ureteral stone (group 2) and ureteral stricture (group 1)**
| **NGAL** | **P value** |
| --- | --- |
| Stone H1 vs. Stricture H1 | 0.153 |
| Stone H2 vs. Stricture H2 | 0.219 |
| Stone H3 vs. Stricture H3 | 0.452 |
| **αGST** |  |
| Stone H1 vs. Stricture H1 | 0.064 |
| Stone H2 vs. Stricture H2 | **0.000** |
| Stone H3 vs. Stricture H3 | **0.000** |
| **πGST** |  |
| Stone H1 vs. Stricture H1 | 0.051 |
| Stone H2 vs. Stricture H2 | **0.003** |
| Stone H3 vs. Stricture H3 | **0.000** |

Bold value represents statistically significant $p < 0.05$ .
NC, negative control; H0, grade 0 hydronephrosis; H1, grade 1 hydronephrosis; H2, grade 2 hydronephrosis; H3, grade 3 hydronephrosis.

### Changes in urine NGAL, αGST, and πGST levels at different time periods

In the ureter stricture group, uNGAL increased significantly on Day 1 and 14 when compared to the baseline, but there was no significant difference between Days 1 and 14 (Fig 1A). However, we didn’t find any significant increase in u-αGST and u-πGST on Day 14 comparing to the baseline.

**Fig 1.**
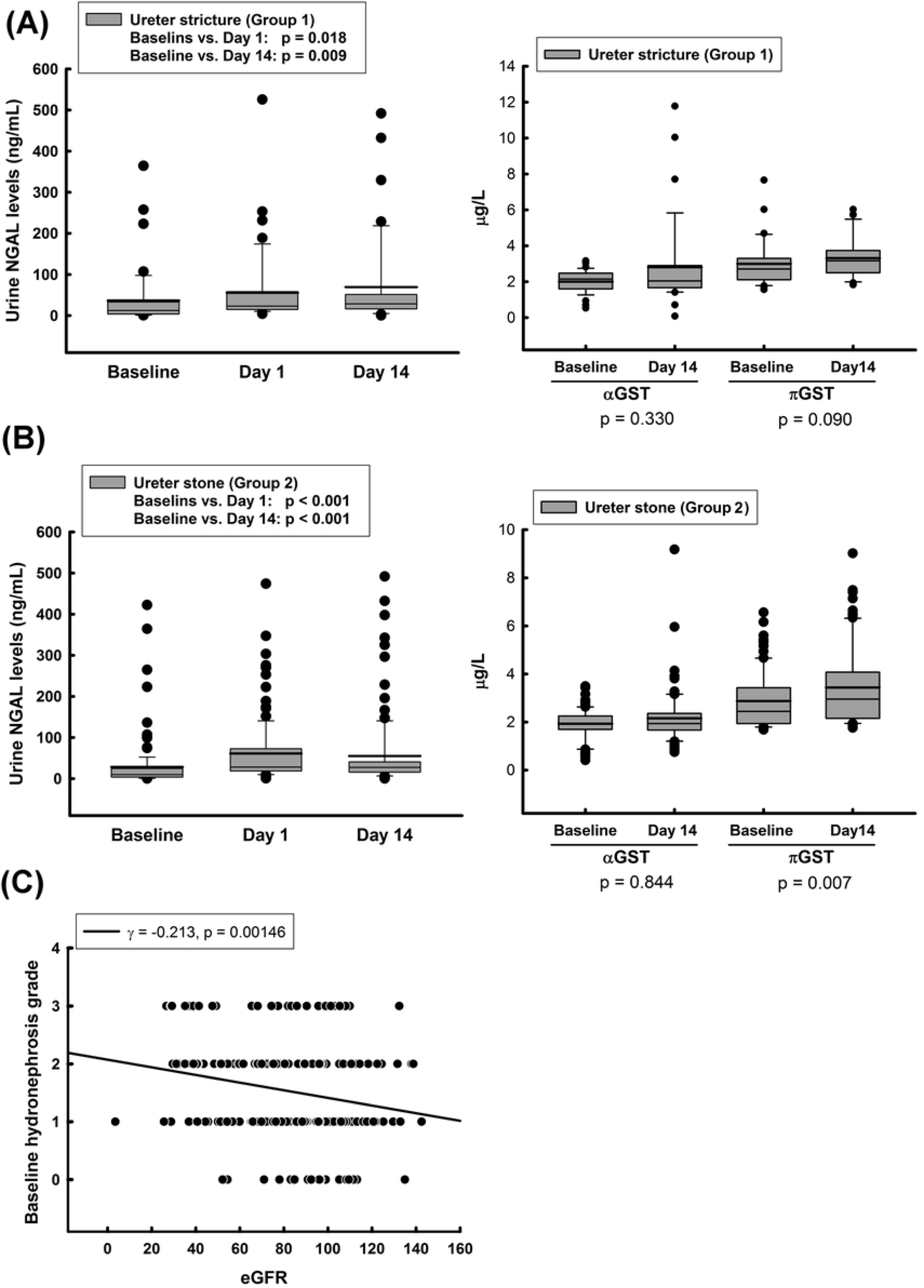
**(A, B)** Changes in urine NGAL, αGST, and πGST levels at baseline, Day 1, and Day 14 after URS in both groups; **(C)** Correlation between baseline hydronephrosis grade and baseline eGFR in all cohorts.

In the ureter stone group, we also found the similar result as the stricture group, that was significant elevated uNGAL on Day 1 and 14 when compared to baseline (Fig 1B). Most importantly, no matter in the group 1 and group 2, there was no significant decrease in the uNGAL from Day 1 to 14, which indicated the injury of kidney related to URS surgery persisted for at least two weeks. On the other hand, u-πGST level increased significantly on day 14 in the stricture group, but the elevation was not noted in u-αGST.

Larger stone may cause more severe obstruction of kidney and more severe kidney injury. Hence, we investigated the impact of stone size on the changes of urinary biomarkers (S1 Fig). The change of uNGAL at different time periods was nearly the same in both ureteral stone ≦1 cm and > 1cm subgroups. However, only in the ureteral stone ≦1 cm subgroup, the level of u-πGST was significantly elevated on Day 14 comparing with the baseline, and no difference between the baseline and Day 14 was found in u-αGST. We further compared three urinary biomarkers between the ureteral stone ≦ 1 cm and > 1cm subgroups. The baseline levels of uNGAL and u-αGST revealed significant difference between stone size ≦ 1 cm and > 1cm subgroups, and the level was significantly higher in the larger stone size group (S1 Fig). However, there was no significant difference in the baseline level of u-πGST but only a non-significant trend (*p*=0.085) (S1 Fig 1).

By Pearson correlation analysis, we found that baseline eGFR and baseline hydronephrosis grade had an inverse correlation significantly (γ = −0.213, p = 0.001) (Fig 1C).

### Predictors of complete recovery of hydronephrosis after URS

We conducted univariate and multivariate logistic regression analysis to find the predictors of complete recovery of hydronephrosis after URS (Table 3). We found that only three factors, including age (OR = 0.96, *p* = 0.002), baseline eGFR (OR = 1.04, *p* < 0.001), and stone size > 1.0 cm (OR = 2.56, *p* = 0.024), had a statistically significant effect on complete recovery of hydronephrosis after URS and after DBJ removal.

**Table 3.** Univariate and multivariate logistic regression analysis for the predictors of complete recovery of hydronephrosis (hydronephrosis grade = 0 on Day 14 after DBJ removal) after URS.

|  | Univariate |  | Multivariate |  |
| --- | --- | --- | --- | --- |
|  | OR (95% CI) | *p* value | OR (95% CI) | *p* value |
| Age , years | 0.97 (0.95, 0.99) | 0.005* | 0.96 (0.94, 0.99) | 0.002* |
| Female (vs Male) | 0.76 (0.42, 1.36) | 0.354 |  |  |
| Group (vs ureteral stricture) |  |  |  |  |
| Ureteral stone | 2.49 (0.30, 4.77) | 0.246 |  |  |
| Staghorn stone | 2.92 (0.81, 10.5) | 0.102 |  |  |
| Stone position |  |  |  |  |
| Upper | 0.62 (0.18, 2.11) | 0.448 |  |  |
| Middle | 0.70 (0.16, 2.98) | 0.630 |  |  |
| Lower | 0.71 (0.20, 2.53) | 0.596 |  |  |
| eGFR (Baseline) | 1.02 (1.01, 1.03) | 0.004* | 1.04 (1.02, 1.06) | <0.001* |
| BMI (kg/m<sup>2</sup>) |  |  |  |  |
| 18.5-24 (vs ≤ 18.5) | 2.17 (0.40, 11.8) | 0.371 |  |  |
| 24-30 (vs ≤ 18.5) | 1.43 (0.28, 7.38) | 0.670 |  |  |
| >30 (vs ≤ 18.5) | 2.57 (0.42, 15.9) | 0.310 |  |  |
| Stone size |  |  |  |  |
| 0.4-1.0 (vs <0.4) | 2.22 (1.06, 4.66) | 0.035* | 1.90 (0.81, 4.44) | 0.138 |
| >1.0 (vs <0.4) | 2.15 (1.07, 4.30) | 0.031* | 2.56 (1.13, 5.77) | 0.024* |
| Presence of UTI | 0.91 (0.51, 1.63) | 0.742 |  |  |
| NGAL(Baseline) | 1.00 (1.00, 1.00) | 0.357 |  |  |
| πGST (Baseline) | 1.06 (0.77, 1.45) | 0.744 | | |
| αGST (Baseline) | 0.83 (0.42, 1.66) | 0.599 | | |
| Cr (Baseline) | 1.02 (0.99, 1.04) | 0.203 |  |  |

Moreover, we investigated the effects of different degrees of hydronephrosis on kidney injury and renal function (Table 4). Each group was stratified into H0+H1 and H2+H3 subgroups. There was no significant difference in eGFR between the two subgroups on baseline and Day 14 in each group. It is worthy to mention that no significant difference in uNGAL between all H2+H3 patients and PC on Day 1, which indicated URS surgery may cause comparable kidney injury as PCNL. Interestingly, till Day 14, the uNGAL from H2+H3 ureteral stricture patient was as high as PC.

**Table 4.** Impact of hydronephrosis degree on eGFR, AKI marker, and urinary kidney damage markers in each group.

|  | Ureteral stricture |  | Ureteral stone |  | Positive control |
| --- | --- | --- | --- | --- | --- |
|  | H0+H1 | H2+H3 | H0+H1 | H2+H3 |  |
| **eGFR**<br>(mL/min/1.73 m<sup>2</sup>) |  |  |  |  |  |
| Baseline | 90.7±4.7 | 79.4±6.7 | 82.5±3.1 | 76.7±3.1* | 89.4±6.6 |
| *P* value | 0.192 |  | 0.323 |  |  |
| Day 14 | 92.7±4.3 | 79.6±8.1 | 87.7±3.4 | 81.2±3.3 | 83.3±6.2 |
| *P* value | 0.176 |  | 0.223 |  |  |
| **NGAL**<br>(ng/mL) |  |  |  |  |  |
| Baseline | 13.6±2.8* | 54.0±16.6 | 19.3±4.4* | 36.1±8.8 | 65.6±16.4 |
| *P* value | 0.125 |  | 0.146 |  |  |
| Day 1 | 33.2±10.9* | 72.4±20.3 | 49.1±9.6*† | 71.4±9.6† | 95.7±24.6 |
| *P* value | **0.017** |  | **0.004** |  |  |
| Day 14 | 18.5±3.5* | 105.0±25.6† | 48.4±10.7*† | 60.5±11.4*† | 137.6±36.3† |
| *P* value | **<0.001** |  | 0.150 |  |  |
| **αGST**<br>(μg/L) |  |  |  |  |  |
| Baseline | 2.1±0.2 | 2.0±0.1 | 2.0±0.1 | 1.9±0.1 | 2.2±0.3 |
| *P* value | 0.684 |  | 0.326 |  |  |
| Day 14 | 3.6±0.8 | 2.3±0.3 | 2.0±0.1* | 2.1±0.2* | 2.5±0.1 |
| *P* value | 0.131 |  | 0.714 |  |  |
| **πGST** |  |  |  |  |  |
| (μg/L) |  |  |  |  |  |
| Baseline | 3.0±0.2 | 3.0±0.3 | 2.7±0.2 | 3.0±0.2 | 2.9±0.6 |
| *P* value | 0.490 |  | 0.691 |  |  |
| Day 14 | 3.7±0.3 | 3.1±0.2 | 3.4±0.2<sup>†</sup> | 3.5±0.3 | 3.2±0.7 |
| *P* value | 0.289 |  | 0.476 |  |  |

H0, hydronephrosis grade 0; H1, hydronephrosis grade 1; H2, hydronephrosis grade 2; H3,
hydronephrosis grade 3; NAGL, neutrophil gelatinase-associated lipocalin; GST, glutathione
S-transferase.
*P* value: the statistical result after analyzed by analysis of variance (ANOVA) in each group.
\* $p < 0.05$ when compared with positive control at the same time period.
Bold value represent statistically significant $p < 0.05$ .
<sup>†</sup> $p < 0.05$ when compared with baseline data in the same subgroup.

### The impact of baseline eGFR on kidney injury

We evaluated the changes in different urine biomarkers (Δ) in baseline eGFR ≧ 60 and eGFR < 60 subgroups in both groups (S1 table). The changes were defined as the level of Day 14 minus the level of baseline. Only ΔNGAL of ureter stone group was significantly higher in eGFR ≧ 60 subgroup than eGFR < 60 subgroup. ΔαGST and ΔπGST had no significant difference between these two subgroups in both study groups.

## Discussion

The present study revealed that no matter in patients with ureteral stones or ureteral stricture, URS surgery would induce AKI and the impact persisted for at least 14 days. The impact of

URS-induced kidney injury, which was indicated by elevated uNGAL in the present study, was more prominent in moderate-to-severe hydronephrosis. In ureteral stone patients with normal renal function (eGFR ≥ 60), uNGAL increased apparently 14 days after URS compared with those with poor renal function (eGFR < 60). However, compared with different size of ureter stones, the degree of URS-induced kidney injury was not significantly different.

### URSL and URS balloon dilatation definitely cause AKI

URS is defined as endoscopic visualization of ureter and renal pelvis for diagnostic and/or therapeutic purposes. Indications for URS include both diagnostic and therapeutic interventions for stone disease, strictures, and ureter tumors in different patient groups [20]. URS is thought to be a safe and effective surgery for treatment of ureteral stones, but the optimal duration and indication for post-URS stenting is controversial. Some authors suggested that post-URS DBJ stent be only used in patients who are at an increased risk of complications, such as previous iatrogenic trauma, impacted ureter calculi, ureter perforation, and under some medical conditions such as solitary kidney, pregnancy, and a history of retroperitoneal fibrosis. Most urologists may favor its use for 1– 2 weeks after URS [20, 21]. Our results suggested that even after DBJ indwelling for 2 weeks, URS-induced AKI still persisted no matter in ureteral stone patients or in ureteral stricture patients. Besides, most noteworthy in the present study is that we used negative and positive controls to assist our interpretations of the results. The negative controls represented no obstructive uropathy, and we compared two study groups with negative controls to validate whether these urinary biomarkers do increase under these clinical conditions. Our results actually proved that both ureter stone and stricture patients with higher degree of hydronephrosis had higher levels of all urinary biomarkers. Furthermore, we used PCNL patients as the positive controls because PCNL is known to create a renal tract to assess intra-pelvic renal stones and it certainly cause considerable kidney injury. There are two interesting finding regarding positive controls. First, although we didn’t see any decrease in the baseline eGFR from positive controls, the baseline level of uNGAL already significantly increased in positive controls compared with mild obstructive uropathy patients in both groups. It indicated that the existence of renal stones would lead to progressive renal injury. Second, in the ureter stricture group with moderate-to-severe hydronephrosis, uNGAL nearly doubled 14 days after URS, which was similar as the positive controls. The level of uNGAL on day 14 in the ureter stricture group with moderate-to-severe hydronephrosis was over 100 ng/mL, even higher than the level on day 1 in positive controls. This finding was not noted In other patients, whereas uNGAL decreased 14 days after a peak in post-URS day 1 in other patients. The possible explanation of this result may be that patients with moderate-to-severe hydronephrosis usually have more stricture and tortuous ureters. The higher severity of ureter stricture definitely increases the difficulty and operation time of URS. However, there was no obvious influence on renal tubule damage after URS except that u-πGST significantly increased in ureteral stone patients with mild hydronephrosis.

### URS procedure causes AKI and tubular damage

Irrigation during URS increases renal pelvic pressure (RPP), possibly leading to intrarenal, pyelo-venous, and pyelo-lymphatic backflow as well as kidney injury [9]. Elevated RPP is noted to be harmful to the kidney in a mini-pig experimental model, and Schwalb et al found that high-pressure irrigation during URS caused irreversible, harmful effects in the kidney, and even moving the URS in the ureter without any irrigation could increase RPP by 20-25 mmHg [22]. Long-term consequence of high RPP (>200 cm H_2_O) caused by high-pressure irrigation is related to columnar metaplasia, subepithelial nests and pericalyceal vasculitis in calyces as compared with those subjected to low irrigant pressure 4-6 weeks after experiment [22]. Our results are compatible with previous studies, which the URS procedure would induce kidney injury, and the elevation in uNGAL can be found as early as 1 hour after URS surgery [23]. URS-related tubular damage is limited to distal tubule and collecting duct in the ureteral stone patients, and our results are consisted with the finding that excessively high collecting system pressure induced renal cellular injury, as reflected by an increase in urinary N-acetyl-β-D-glucosaminidase levels (non-specific renal tubular marker) [24].

### Ureter Balloon dilation may cause more AKI than Ho:YAG laser lithotripsy

High-grade hydronephrosis (H2+H3) caused more AKI on Day 1 as noted in both study groups, and even more AKI was present on Day 14 in the H2+H3 subgroup of ureter stricture group (Table 4). This result implied that URS balloon dilation may cause more AKI than URSL in the high-grade hydronephrosis patients on Day 14. Acute high RPP (>200 cmH_2_O) not only causes diffuse denudation and flattening of the caliceal urothelium, submucosal edema and congestion [22], but also causes renal tubule dilation and renal glomerulus compression [25]. This may be the reason why the elevated uNGAL levels in the ureteral stricture group were as high as those in the PCNL group (PC), known to have a sudden and rapid increase in RPP and direct renal parenchymal injury during the procedure. On the contrary, the mechanism of Ho:YAG laser lithotripsy is through photothermal effects with minimal upward migration of ureteral stone during procedure. Though intermittent normal saline irrigation by hand-held syringe during Ho:YAG laser lithotripsy, there will be no renal injury when the RPP was maintained under 120 cmH_2_O (around 88 mmHg) [22]. In a recent porcine experimental study, under gravity irrigation and manual pumping, the maximal RPP during URS reached 30 and 105 cmH_2_O, respectively, which were all lower than 120 cnH_2_O [8]. Therefore, in our current study, the uNGAL level on Day 14 in ureteral stone group was significantly lower than both the ureteral stricture group and PCNL group. Besides, only the H2+H3 subgroup of ureter stricture group had persistent elevated uNGAL levels on Day 1 to Day 14, which was also seen in PCNL group patients. This interesting finding suggests the effect of URS produce for severe ureteral stricture complicated with high-grade hydronephrosis may cause prolonged kidney injury.

### The severity of hydronephrosis affects post-URS AKI, but not tubule damage

The kidney is the organ with highest blood flow in the human body, which receives approximately one fourth of the cardiac output. Obstructive nephropathy refers to anatomical or functional obstruction of the kidney and leads to progressive kidney injury. Using CT perfusion image, Cai et al. found that significantly decrease in blood flow in both renal cortex and medulla from the obstructive kidney of moderate and severe unilateral ureteral obstruction (UUO); whereas there was no significantly difference in mean blood flow, blood volume, and clearance in the obstructive kidney of mild UUO [26]. This finding suggested that mild dilation of pelvis and calyx is not enough to deteriorate the renal function. This is why H0 and H1 hydronephrosis in both groups 1 and 2 had no significant uNGAL elevation after URS from baseline to Day 14, and the patients with H2 and H3 hydronephrosis had the same severity of uNGAL elevation on Day 1 when compared to the PCNL group. However, mean u-αGST and u-πGST levels were not influenced by URS surgery in both study groups, which implied these three urinary cytokines represented the different types of kidney injury.

## Conclusions

Although eGFR didn’t significantly change after URS surgery, uNGAL persisted elevation for two weeks in both ureteral stricture and ureteral stone groups, which suggested that URS procedures could cause kidney injury. With respect to renal tubular damage marker, only u-πGST was elevated on Day 14 in the ureteral stone group. Ureteral stone patients with preserved renal function suffered more uNGAL changes (i.e. ΔNGAL) after URS. Taken together, both URSL and URS balloon dilation could lead to kidney injury and renal distal tubule damage till two weeks even though DBJ indwelling continued in both groups.

## References

1. Ma MC, Huang HS, Chen CF. Impaired renal sensory responses after unilateral ureteral obstruction in the rat. J Am Soc Nephrol. 2002;13(4):1008–16.

2. Harris RH, Yarger WE. Renal function after release of unilateral ureteral obstruction in rats. Am J Physiol. 1974;227(4):806–15.

3. Ulrich JC, York JP, Koff SA. The renal vascular response to acutely elevated intrapelvic pressure: resistive index measurements in experimental urinary obstruction. J Urol. 1995;154(3):1202–4.

4. Tsujihata M. Mechanism of calcium oxalate renal stone formation and renal tubular cell injury. Int J Urol. 2008;15(2):115–20.

5. Duan X, Kong Z, Mai X, Lan Y, Liu Y, Yang Z, et al. Autophagy inhibition attenuates hyperoxaluria-induced renal tubular oxidative injury and calcium oxalate crystal depositions in the rat kidney. Redox Biol. 2018;16:414–25.

6. Huang HS, Liao PC, Liu CJ. Calcium Kidney Stones are Associated with Increased Risk of Carotid Atherosclerosis: The Link between Urinary Stone Risks, Carotid Intima-Media Thickness, and Oxidative Stress Markers. J Clin Med. 2020;9(3).

7. Hofmann R. [Ureteroscopy (URS) for ureteric calculi]. Urologe A. 2006;45(5):W637–46; quiz W47.

8. Noureldin YA, Kallidonis P, Ntasiotis P, Adamou C, Zazas E, Liatsikos EN. The Effect of Irrigation Power and Ureteral Access Sheath Diameter on the Maximal Intra-Pelvic Pressure During Ureteroscopy: In Vivo Experimental Study in a Live Anesthetized Pig. J Endourol. 2019;33(9):725–9.

9. Tokas T, Herrmann TRW, Skolarikos A, Nagele U, Training, Research in Urological S, et al. Pressure matters: intrarenal pressures during normal and pathological conditions, and impact of increased values to renal physiology. World J Urol. 2019;37(1):125–31.

10. Tokas T, Skolarikos A, Herrmann TRW, Nagele U, Training, Research in Urological S, et al. Pressure matters 2: intrarenal pressure ranges during upper-tract endourological procedures. World J Urol. 2019;37(1):133–42.

11. Gnessin E, Yossepowitch O, Holland R, Livne PM, Lifshitz DA. Holmium laser endoureterotomy for benign ureteral stricture: a single center experience. J Urol. 2009;182(6):2775–9.

12. Washino S, Hosohata K, Miyagawa T. Roles Played by Biomarkers of Kidney Injury in Patients with Upper Urinary Tract Obstruction. Int J Mol Sci. 2020;21(15).

13. Kuntz NJ, Neisius A, Tsivian M, Ghaffar M, Patel N, Ferrandino MN, et al. Balloon Dilation of the Ureter: A Contemporary Review of Outcomes and Complications. J Urol. 2015;194(2):413–7.

14. Siew ED, Ware LB, Gebretsadik T, Shintani A, Moons KG, Wickersham N, et al. Urine neutrophil gelatinase-associated lipocalin moderately predicts acute kidney injury in critically ill adults. J Am Soc Nephrol. 2009;20(8):1823–32.

15. Schmidt-Ott KM, Mori K, Li JY, Kalandadze A, Cohen DJ, Devarajan P, et al. Dual action of neutrophil gelatinase-associated lipocalin. J Am Soc Nephrol. 2007;18(2):407–13.

16. Yang HN, Boo CS, Kim MG, Jo SK, Cho WY, Kim HK. Urine neutrophil gelatinase-associated lipocalin: an independent predictor of adverse outcomes in acute kidney injury. Am J Nephrol. 2010;31(6):501–9.

17. Huang HS, Ma MC. High Sodium-Induced Oxidative Stress and Poor Anticrystallization Defense Aggravate Calcium Oxalate Crystal Formation in Rat Hyperoxaluric Kidneys. PLoS One. 2015;10(8):e0134764.

18. Ma MC, Chen YS, Huang HS. Erythrocyte oxidative stress in patients with calcium oxalate stones correlates with stone size and renal tubular damage. Urology. 2014;83(2):510 e9–17.

19. Walshe CM, Odejayi F, Ng S, Marsh B. Urinary glutathione S-transferase as an early marker for renal dysfunction in patients admitted to intensive care with sepsis. Crit Care Resusc. 2009;11(3):204–9.

20. Whitehurst LA, Somani BK. Semi-rigid ureteroscopy: indications, tips, and tricks. Urolithiasis. 2018;46(1):39–45.

21. Somani BK, Giusti G, Sun Y, Osther PJ, Frank M, De Sio M, et al. Complications associated with ureterorenoscopy (URS) related to treatment of urolithiasis: the Clinical Research Office of Endourological Society URS Global study. World J Urol. 2017;35(4):675–81.

22. Schwalb DM, Eshghi M, Davidian M, Franco I. Morphological and physiological changes in the urinary tract associated with ureteral dilation and ureteropyeloscopy: an experimental study. J Urol. 1993;149(6):1576–85.

23. Benli E, Ayyildiz SN, Cirrik S, Noyan T, Ayyildiz A, Cirakoglu A. Early term effect of ureterorenoscopy (URS) on the Kidney: research measuring NGAL, KIM-1, FABP and CYS C levels in urine. Int Braz J Urol. 2017;43(5):887–95.

24. Fung LC, Atala A. Constant elevation in renal pelvic pressure induces an increase in urinary N-acetyl-beta-D-glucosaminidase in a nonobstructive porcine model. J Urol. 1998;159(1):212–6.

25. Wang J, Zhou DQ, He M, Li WG, Pang X, Yu XX, et al. Effects of renal pelvic high-pressure perfusion on nephrons in a porcine pyonephrosis model. Exp Ther Med. 2013;5(5):1389–92.

26. Cai XR, Zhou QC, Yu J, Feng YZ, Xian ZH, Yang WC, et al. Assessment of renal function in patients with unilateral ureteral obstruction using whole-organ perfusion imaging with 320-detector row computed tomography. PLoS One. 2015;10(4):e0122454.

